# Establishing live-cell single-molecule localization microscopy imaging and single-particle tracking in the archaeon *Haloferax volcanii*

**DOI:** 10.1101/2020.07.27.222935

**Authors:** Bartosz Turkowyd, Sandra Schreiber, Julia Wörtz, Ella Shtifman Segal, Moshe Mevarech, Iain G. Duggin, Anita Marchfelder, Ulrike Endesfelder

**Affiliations:** Department of Systems and Synthetic Microbiology, Max Planck Institute for Terrestrial Microbiology and LOEWE Center for Synthetic Microbiology (SYNMIKRO), 35043 Marburg, Germany; Department of Biology II, Ulm University, 89069 Ulm, Germany; Department of Molecular Microbiology and Biotechnology, George S. Wise Faculty of Life Sciences, Tel Aviv University, Tel Aviv 69978-01, Israel; The ithree institute, University of Technology Sydney, Ultimo, NSW, 2007, Australia; Department of Physics, Mellon College of Science, Carnegie-Mellon University, Pittsburgh, PA 15213, USA

**Keywords:** single-molecule imaging and tracking, advanced fluorescence microscopy, archaeal cell imaging, imaging technology in microbiology, Haloferax volcanii, FtsZ1 division ring, RNA polymerase dynamics

## Abstract

In recent years, fluorescence microscopy techniques for the localization and tracking of single molecules in living cells have become well-established and indispensable tools for the investigation of cellular biology and *in vivo* biochemistry of many bacterial and eukaryotic organisms. Nevertheless, these techniques are still not established for imaging archaea. Their establishment as a standard tool for the study of archaea will be a decisive milestone for the exploration of this branch of life and its unique biology.

Here we have developed a reliable protocol for the study of the archaeon *Haloferax volcanii*. We have generated an autofluorescence-free *H. volcanii* strain, evaluated several fluorescent proteins for their suitability to serve as single-molecule fluorescence markers and codon-optimized them to work under optimal *H. volcanii* cultivation conditions. We found that two of them, Dendra2Hfx and PAmCherry1Hfx, provide state-of-the-art single-molecule imaging. Our strategy is quantitative and allows dual-color imaging of two targets in the same field of view as well as DNA co-staining. We present the first single-molecule localization microscopy (SMLM) images of the subcellular organization and dynamics of two crucial intracellular proteins in living *H. volcanii* cells, FtsZ1, which shows complex structures in the cell division ring, and RNA polymerase, which localizes around the periphery of the cellular DNA. This work should provide incentive to develop SMLM strategies for other archaeal organisms in the near future.

## Introduction

The development of super-resolution fluorescence microscopy techniques that can achieve resolutions in the nanometer range, and thus enable the investigation of sub-cellular structures in living cells, has considerably expanded the possibilities of cell biology in the life sciences (Turkowyd et al., 2016;Sigal et al., 2018). These new techniques include Single-Molecule Localization Microscopy (SMLM), which has the unique ability to study individual molecules and measure their actions in live cells with high specificity, sensitivity and spatiotemporal resolution. While single molecules could previously only be studied in the controlled, but limited environment of *in vitro* assays, SMLM imaging allows us to measure their function in their native cellular environment. Consequently, SMLM imaging has become a powerful tool for studying cell biology (Endesfelder, 2019).

SMLM imaging has become a well-established technique for many bacteria and yeast in recent years (Vojnovic et al., 2019), but archaea have not yet been studied using this technique. For archaeal studies, even standard fluorescence microscopy is still only a rarely used tool, as fluorescence imaging faces a) the often extreme growth conditions of archaeal organisms (e.g. very high or low temperature or high saline, acidic, alkaline or anaerobic environments), b) their colorful pigments causing autofluorescence and c) the limited range of available model organisms and molecular tools (Kramm et al., 2019).

The set of fluorescence tools for imaging archaea available today includes several optimized fluorescent protein versions, a codon-optimized and a split version of GFP (Reuter and Maupin-Furlow, 2004;Duggin et al., 2015;Winter et al., 2018;Li et al., 2019) and a mCherry (Duggin et al., 2015;Liao et al., 2020) for imaging of Haloarchaea and a heat-tolerant GFP for *Sulfolobus* (Pulschen et al., 2020). Extrinsic stains include DNA staining with Acridine Orange, propidium iodide, Hoechst 33258 or 33342, DAPI and SYTOX Green (Samson et al., 2008, Liao, 2020 #2010;Delmas et al., 2013) or BrdU and EdU labeling of nascent DNA (Gristwood et al., 2012;Delpech et al., 2018), FM4-64Xa staining of membranes (Samson et al., 2008) or non-permeable dyes that stain the cell wall (Wirth et al., 2011), a new thermostable protein tag (H5) for *Sulfolobus* (Visone et al., 2017) and some immunofluorescence staining of fixed cells (Poplawski et al., 2000;Lindas et al., 2008;Samson et al., 2008;Ettema et al., 2011;Zenke et al., 2015).

Establishing advanced fluorescence microscopy as a regular instrument for the study of archaea will be a decisive milestone for the exploration of this branch of life (Bisson-Filho et al., 2018). Archaea encompass a mixture of bacterial, eukaryotic and unique characteristics (Leigh et al., 2011), thus the study of the cellular functions of archaeal proteins will reveal new and important aspects of biology. In addition, due to their unusual, extreme habitats, archaea promise the discovery of a highly interesting and surprising molecular biology not found elsewhere.

In this report we present the first protocol for the investigation of *Haloferax volcanii* using SMLM methods. We have developed a background-reduced *H. volcanii* strain by interrupting its carotenoid synthesis pathway, evaluated and codon-optimized suitable fluorescent proteins as genetic fluorescent markers and established reliable sample preparation protocols under high salt and temperature conditions in different media. We demonstrate that our SMLM readout is both quantitative and allows for dual-color SMLM imaging and DNA co-staining. We present the first SMLM images of two crucial intracellular proteins in live *H. volcanii*, namely the sub-cellular architecture of the division ring protein FtsZ1 and the spatiotemporal distribution of the RNA polymerase.

## Materials & Methods

### Generation of *Haloferax* strain WR806

For generating the deletion strain WR806 the plasmid pMM1260 was constructed, which contains the flanking regions of *crtI* (HVO_2528) (500 bp upstream and downstream of that gene). The pop-in/pop-out method (Bitan-Banin et al., 2003) was applied for transformation using the PEG600 protocol as described in the HaloHandbook (Dyall-Smith, 2009). *Haloferax* H133 cells (Allers et al., 2004) were transformed with pMM1260 to mediate integration of the plasmid into the *crtl* gene. Transformed colonies were then selected for by plating on Ca medium supplemented with tryptophan and thymidine. Pop-in candidates were plated on medium containing 5-FOA (5-Fluoroorotic acid) to select for pop-out clones. Pop-out strains were verified using PCR and selective plating.

### Plasmids expressing fluorescence proteins (pTA231)

*E. coli* codon-optimized genes encoding PAmCherry1 and mMaple3 (Wang et al., 2014) were amplified using Phusion polymerase from plasmids pPAmCherry-CAM and pmMaple3-CAM, respectively, by using oligonucleotides providing *Nde*I*/Xba*I restriction sites. PCR products were first cloned into pBluescript via blunt DNA ends yielding plasmids pBlue-PAmCherry1opt and pBlue-linker-mMaple3opt. Genes were excised from these plasmids with *Nde*I and *Xba*I and subsequently ligated into plasmid pTA231-p.Syn (digested with *Nde*I and *Xba*I) yielding plasmids pTA231-p.Syn-PAmCherry1 and pTA231-p.Syn-mMaple3. Plasmid pDendra2-CAM (Wang et al., 2014) was used as a template for amplification of Dendra2, with oligonucleotides containing *Xho*I and *Xba*I restriction sites. The Phusion PCR product was first cloned into pBluescript via blunt DNA ends yielding plasmid pBlue-Dendra2. The Dendra2 gene was obtained by digestion of this plasmid with *Xho*I, the resulting 5’-overhang was converted to double stranded DNA by treatment with DNA polymerase I Klenow fragment. The linearized plasmid was then digested with *Xba*I releasing the fragment containing the gene. Plasmid pTA231-p.Syn was digested with *Nde*I, followed by DNA polymerase I Klenow fragment treatment, before *Xba*I digestion of the linearized plasmid. Finally, the DNA fragment Dendra2 was ligated with the digested pTA231-p.Syn resulting in plasmid pTA231-p.Syn-Dendra2.

Dendra2 and PAmCherry1 genes were obtained from GeneArt® (Thermo Fischer Scientific, Waltham, USA) as codon-optimized versions for expression in *H. volcanii* (plasmids pMA-RQ-Dendra2Hfx and pMA-T-PAmCherry1Hfx). The Dendra2Hfx gene was obtained from pMA-RQ-Dendra2Hfx by digestion with *Nde*I and *Xba*I, the resulting fragment was ligated into pTA231-p.Syn (digested with *Nde*I and *Xba*I) yielding plasmid pTA231-p.Syn-Dendra2Hfx. The PAmCherry1Hfx gene was amplified by PCR using oligonucleotides providing *Nde*I/*Xba*I restriction sites and pMA-T-PAmCherry1Hfx as template. After cloning into pBluescript via blunt DNA ends (yielding pBlue-PAmCherry1Hfx), the gene was excised with *Nde*I*/Xba*I and cloned into pTA231-p.Syn, yielding plasmid pTA231-p.Syn-PAmCherry1Hfx. *H. volcanii* WR806 cells were transformed with these plasmids and grown in Hv-Cab medium (Duggin et al., 2015) with 0.45 mM uracil and 0.16 mM thymidine.

### Plasmids expressing fluorescence-fusion proteins (pTA962)

The fluorescence fusion constructs used in this study were based on the plasmid pTA962-FtsZ1-smRSGFP (also termed pIDJL40-FtsZ1) (Duggin et al., 2015). To obtain plasmids pTA962-FtsZ1-Dendra2Hfx and pTA962-FtsZ1-PAmCherry1Hfx, the smRSGFP sequence from pTA962-FtsZ1-smRSGFP was excised with *Not*I/*Bam*HI and the resulting plasmid was ligated with genes for Dendra2Hfx or PAmCherry1Hfx, that were obtained by amplification from pMA-RQ-Dendra2Hfx using oligonucleotides containing suitable restriction sites and digestion of pMA-T-PAmCherry1Hfx, respectively. To obtain pTA962-RpoD-Dendra2Hfx, the *rpoD* gene (HVO_2781) was amplified from H119 genomic DNA via Phusion PCR using primers providing *Nde*I*/Bam*HI restriction sites, the resulting fragment was cloned via blunt ends into pBluescript (digested with *Eco*RV), yielding pBlue-RpoD. The *rpoD* gene was excised from pBlue-RpoD using *Nde*I/*Bam*HI. Plasmid pTA962-FtsZ1-Dendra2Hfx was digested with *Nde*I/*Bam*HI (which releases the *ftsz1* gene) and the *rpoD* gene was ligated to the linearized plasmid, yielding pTA962-RpoD-Dendra2Hfx. *H. volcanii* WR806 cells were transformed with these plasmids and grown in Hv-Cab medium with 0.25 mM tryptophan (defined medium) or YPC (full medium).

### Preparation of cultures for fluorescence microscopy

All liquid cultures were prepared from cryo stocks in Hv-Cab media unless stated otherwise. Cultures of strains expressing pTA231 were additionally supplied with 0.45 mM uracil and 0.16 mM thymidine, and cultures of strains expressing pTA962 were supplied with additional 0.25 mM tryptophan. WR806 and H119 cultures not expressing either pTA231 or pTA962 were supplied with 0.45 mM uracil, 0.16 mM thymidine and 0.25 mM tryptophan.

After inoculation, cultures were incubated for 72 hours at 42°C shaking at 200 RPM and diluted with fresh Hv-Cab media containing appropriate nucleobases and amino acids to a final OD600 of 0.1. WR806 strains expressing pTA962-FtsZ1-Dendra2Hfx and pTA962-RpoD-Dendra2Hfx were additionally cultured in YPC media for comparative studies shown in figures 3 and 4. After inoculation, YPC cultures were incubated for 48 hours at 42°C shaking at 200 RPM and diluted afterwards with fresh YPC media to a final OD600 of 0.1. Diluted Hv-Cab and YPC cultures were incubated for another three hours at 42°C shaking at 200 RPM. 750 µL of each culture was centrifuged (5.000 x g, 3 minutes), supernatants were discarded and pellets were suspended in 50 µL of fresh Hv-Cab media. At this step, Hoechst 33342 was added to cell suspensions to a final concentration of 10 µg/mL for 10 minutes, and suspensions were washed once with fresh Hv-Cab media to remove the excess of Hoechst 33342. 2 µL of suspensions were placed on agarose pads and covered with a cleaned coverslip.

**Figure 1.**
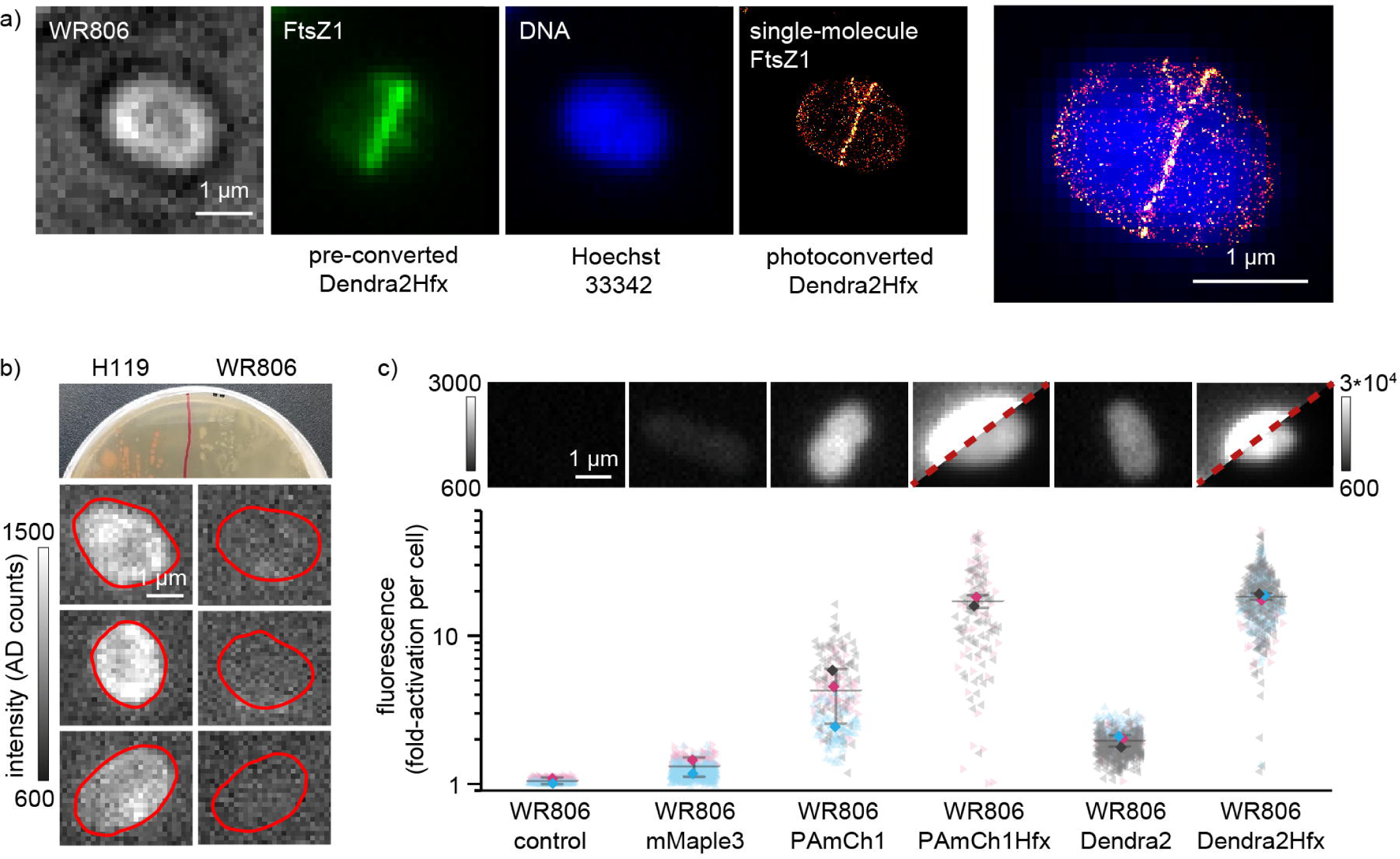
Establishing SMLM imaging in *Haloferax volcanii*. a) A representative image of a WR806 cell expressing the fluorescently tagged FtsZ1. The fluorescent protein Dendra2Hfx can be read-out in a diffraction-limited mode using its pre-converted green fluorescent form as well as in a single-molecule readout mode using photoconverted FtsZ1-Dendra2Hfx. This provides a high-resolution and quantitative SMLM image of FtsZ1 that reveals fine structural details. Diffraction limited snapshots of DNA-Hoechst 33342 were taken after SMLM imaging. b) The H119 strain synthesizes lycopene and its derivatives (Haque et al., 2020) which can be easily seen by bright red colonies on agar-plates. These cause autofluorescence background that largely hinders single-molecule readout. The WR806 strain lacks the phytoene dehydrogenase *crtI* of the lycopene synthesis pathway, which results in white cells on plates and drastically reduces autofluorescence. c) Three SMLM-suitable fluorescent proteins (mMaple3, PAmCherry1 and Dendra2) were evaluated for their expression and functionality in the cytosol of *H. volcanii* cells. Their fluorescence signal before and after photoactivation/-conversion was measured. mMaple3 showed no substantial signal (readout levels similar to a non-fluorescent WR806 control), PAmCherry1 and Dendra2 gave a fluorescent signal after photoactivation/-conversion. After codon-optimization, Dendra2Hfx and PAmCherry1Hfx showed an increase in signal by one order of magnitude. Experiments were done in two or three independent replicates which are color-coded in red, blue and dark grey. The mean of each replicate is marked by a diamond shape. The overall means are marked by horizontal lines. Error bars represent standard deviations of replicate means. Statistics: WR806 control 164 cells (two replicates); WR806 mMaple3 474 cells (two replicates); WR806 PamCherry11 226 cells (three replicates); WR806 PAmCherry1Hfx 129 cells (two replicates); WR806 Dendra2 566 cells (three replicates); WR806 Dendra2Hfx 475 cells (three replicates). The diagonal red dashed line for cells expressing codon-optimized Dendra2Hfx and PAmCherry1Hfx separates two representations with different dynamic range. The left part of the image has the same dynamic range used for all exemplary images (as given by the scale to the left), the right of the image a ten-times wider dynamic range (as given by the scale to the right).

**Figure 2.**
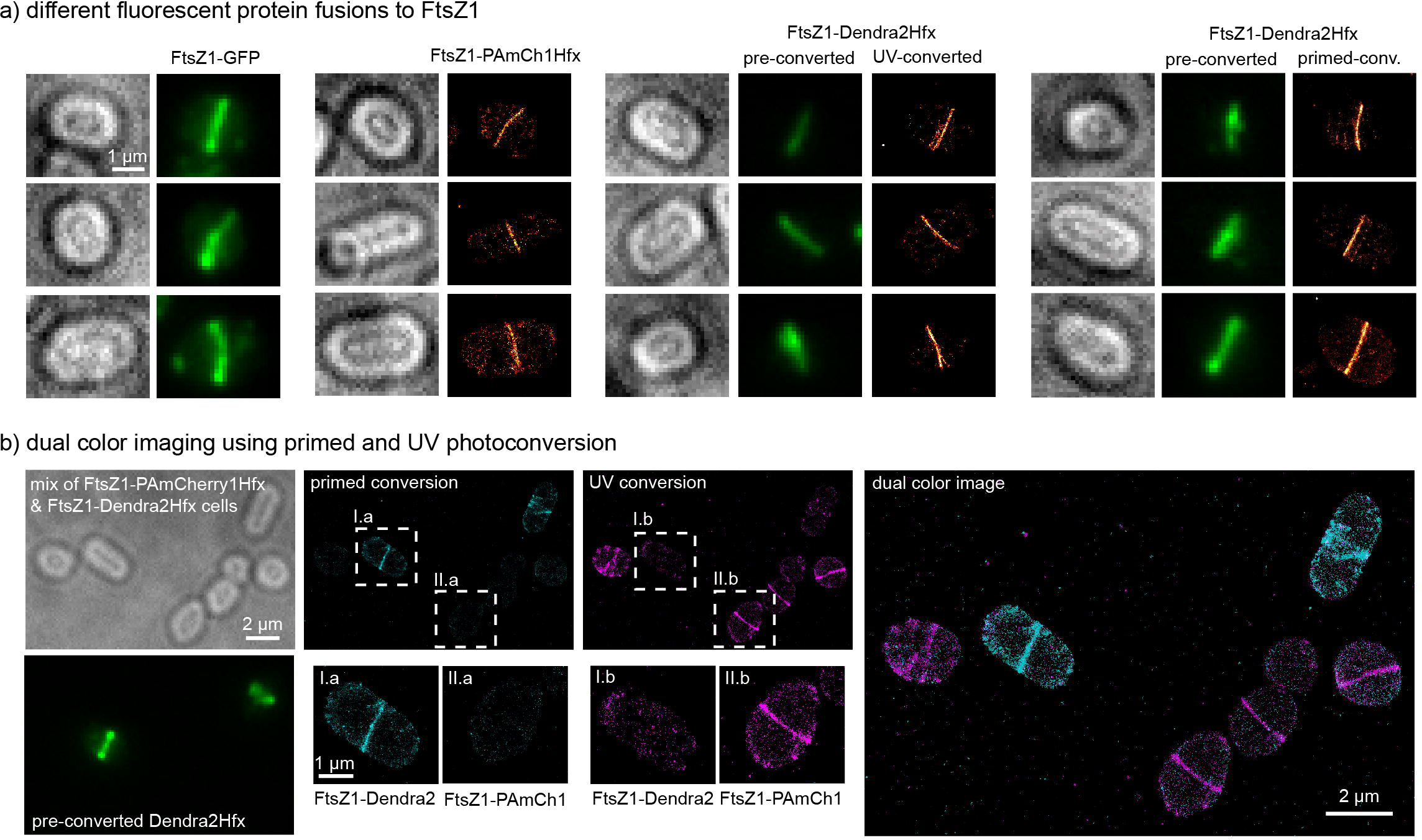
Fluorescent imaging of PamCherry1Hfx and Dendra2Hfx-labeled FtsZ1 proteins in *Haloferax volcanii*. a) Cells expressing FtsZ1-GFP show a distinct FtsZ1 ring structure at mid cell (green signal). By using PAmCherry1Hfx and Dendra2Hfx, the FtsZ1 ring can be imaged with higher spatial resolution providing fine structural details. FtsZ1-PAmCherry1Hfx and FtsZ1-Dendra2Hfx are efficiently photoactivated/-converted using 405 nm light. In addition, FtsZ1-Dendra2Hfx can be photoconverted using primed photoconversion with 488 nm and 730 nm light. The experimental single-molecule localization precisions that were obtained were measured by the NeNA approach (Endesfelder et al., 2014) and amounted to 12 nm. b) A composite sample was prepared by mixing WR806 cells expressing either FtsZ1-Dendra2Hfx or FtsZ1-PAmCherry1Hfx. Strains can be differentiated in the green channel as PAmCherry1Hfx lacks fluorescence before photoactivation. First, FtsZ1-Dendra2Hfx was readout via primed photoconversion (I.a), while FtsZ1-PAmCherry1Hfx remained dark (II.a). After full read-out of FtsZ1-Dendra2Hfx (I.b), FtsZ1-PAmCherry1Hfx was readout by UV-photoactivation (II.b).

**Figure 3.**
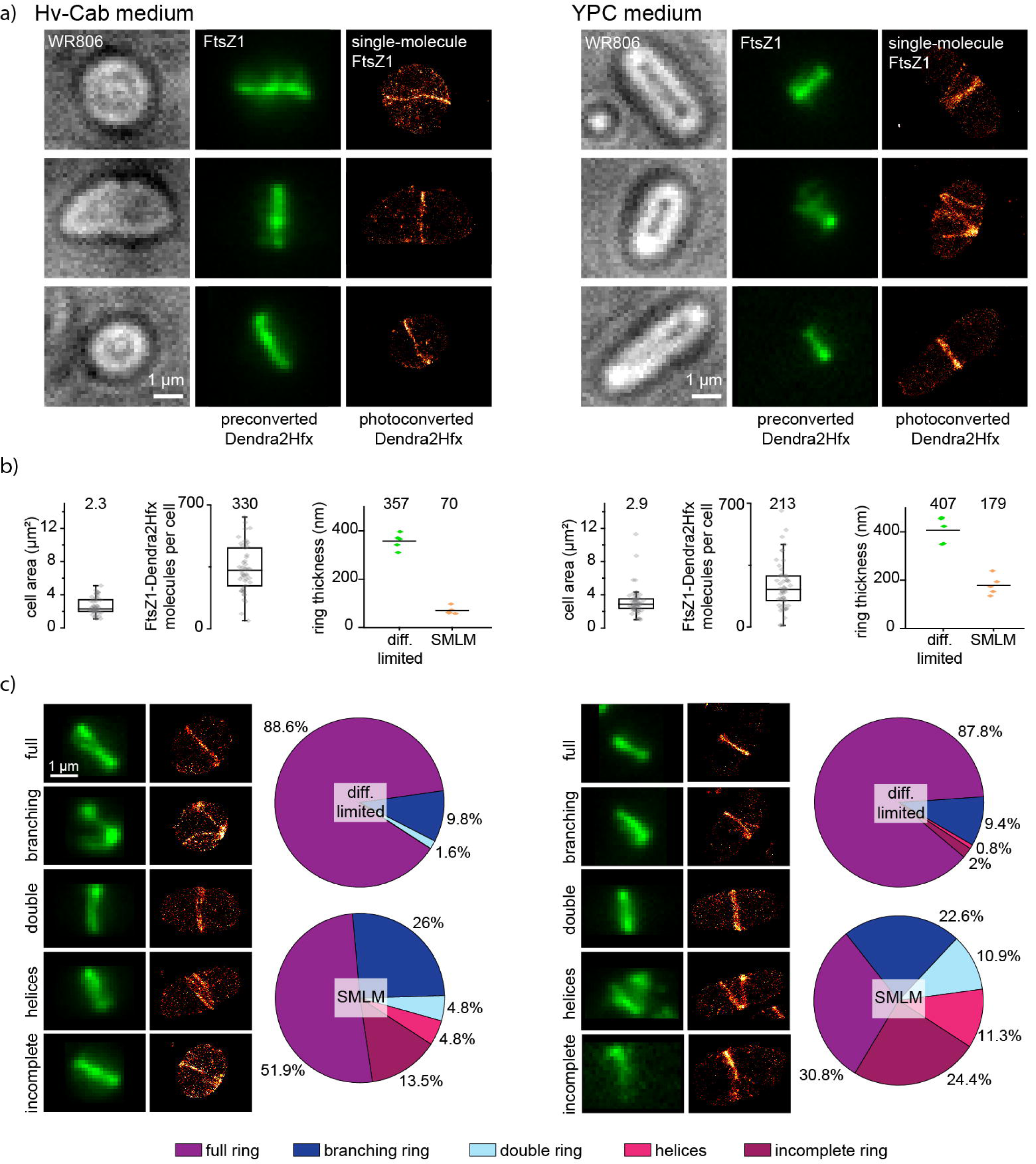
Resolving the FtsZ1-Dendra2Hfx ring structure in *H. volcanii*. a) Representative images of FtsZ1 structures in Hv-Cab and YPC media. Epifluorescence images of pre-converted Dendra2Hfx did not show any substantial differences in ring structures for different growth conditions, whereas SMLM imaging of photoconverted Dendra2Hfx revealed highly diverse ring structures. The experimental single-molecule localization precisions that were obtained were measured by the NeNA approach (Endesfelder et al., 2014) and amount to 12 nm (Hv-Cab) and 13 nm (YPC). b) While cellular areas are similar between different growth conditions, cells growing in Hv-Cab possess similar numbers of labeled FtsZ1 protein copies than cells cultivated in YPC media. In addition, their FtsZ1 ring structures are thinner. Boxes represent 25-75% data range, horizontal lines median values and whiskers mark the outlier range with coefficient 1.5. Statistics for the cell area analysis and FtsZ1 protein counting: Hv-Cab 50 cells; YPC 54 cells. Statistics for ring thickness measurements: 5 cells for both growth conditions. c) FtsZ1 structures visualized in epifluorescence and SMLM were assigned to five classes: full, branching, double, helical and incomplete rings. Statistics: Hv-Cab 132 cells; YPC 256 cells.

**Figure 4.**
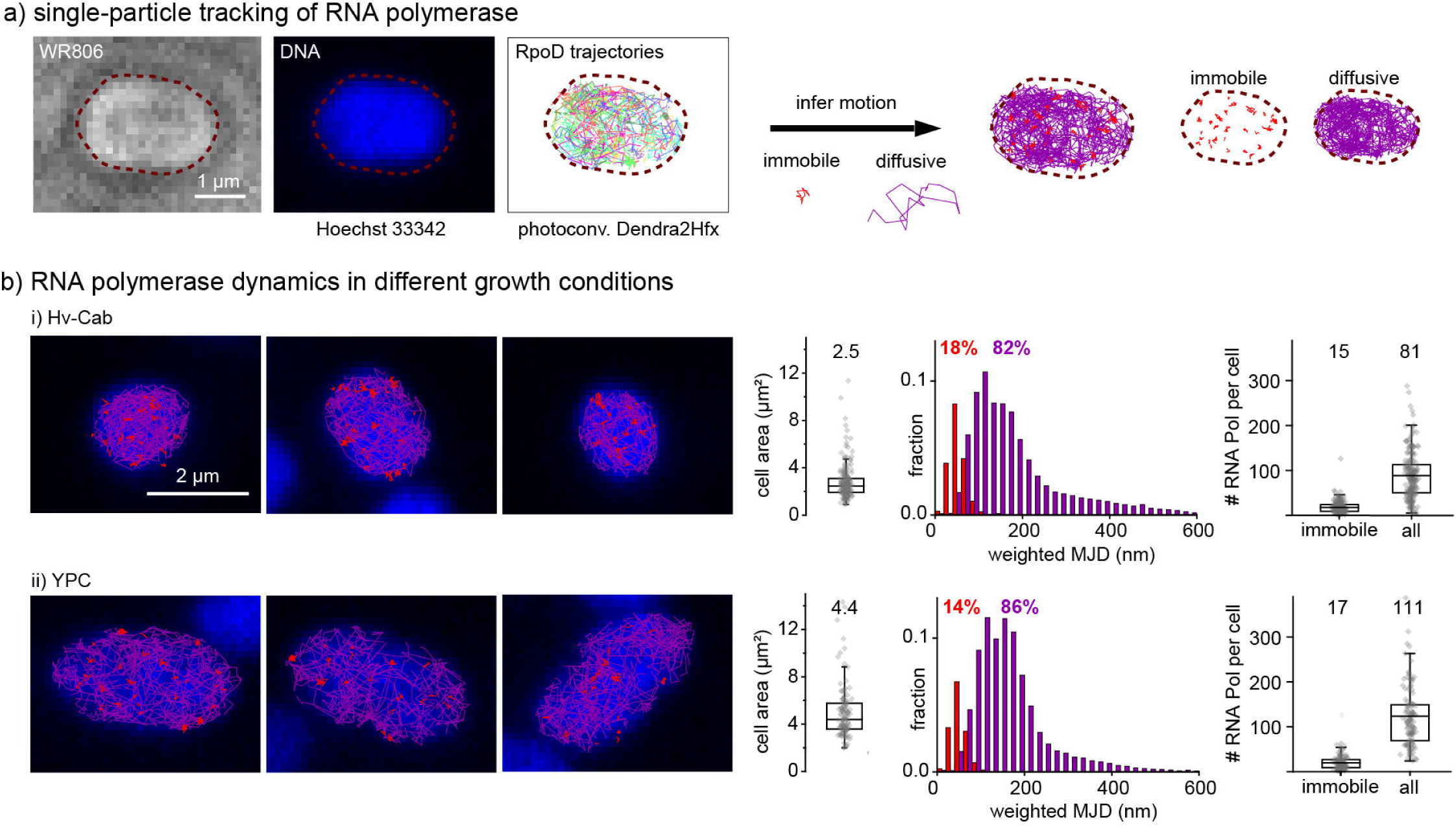
Single-molecule tracking of RNA polymerase in *H. volcanii*. a) WR806 cells expressing RpoD-Dendra2Hfx were imaged by SMLM imaging. The bright light image (left inset) was taken prior to SMLM recording, the DNA image (middle inset) afterwards (to prevent unnecessary photoconversion of Dendra2Hfx). Single-molecule localizations were connected into trajectories and a trajectory map is reconstructed (right inset). Each trajectory was assigned to immobile or diffusive motion (blue and violet, respectively). b) SMLM imaging reveals differences in RNA polymerase dynamics for two different growth conditions: i) Hv-Cab medium optimized for imaging and ii) rich YPC medium. Cells growing in Hv-Cab media are approx. 50% smaller in their area compared to cells growing in the richer YPC medium. Despite different growth conditions there was no significant difference in the number of immobile, actively transcribing RNA polymerases per cell (15 and 17 RNA polymerase copies detected throughout imaging per cell for Hv-Cab and YPC, respectively) while the overall number of labeled RNA polymerase copies was lower in Hv-Cab (81 and 111 copies per cell for Hv-Cab and YPC, respectively). Boxes represent 25-75% data range, horizontal lines median values and whiskers mark the outlier range with coefficient 1.5. Statistics: Hv-Cab 198 cells; YPC 115 cells.

### Agarose pad preparation

1% w/v low melting agarose (Merck/Sigma-Aldrich, Germany) stock was prepared by suspending the agarose powder in 18% BSW (Dyall-Smith, 2009). The mixture was incubated at 70°C for 12 minutes to melt the agarose and then cooled down to 42°C. The agarose solution at 42°C was placed on indented microscope slides (Thermo Fisher, Germany) and sealed with coverslips cleaned overnight in 1 M KOH (Merck/Sigma Aldrich). After three hours, agarose pads were ready to use.

### Single-molecule localization microscopy setup

All SMLM studies were performed on a custom-built setup based on a Nikon Ti Eclipse microscope equipped with a set of dichroic mirrors and filters (ET dapi/Fitc/cy3 dichroic, ZT405/488/561rpc rejection filter, ET525/50 or ET610/75 bandpass, all AHF Analysentechnik, Germany), and a CFI Apo TIRF 100x oil objective (NA 1.49, Nikon) which was heated to 37°C. 405 nm OBIS, 488 nm Sapphire and 561 nm OBIS lasers (all Coherent Inc., USA) were controlled via an acousto-optical tunable filter (AOTF) (Gooch and Housego, USA). The 730 nm OBIS laser (Cohrent Inc., USA) was controlled manually using the manufacturer’s software. Fluorescence was detected by an emCCD (iXON Ultra 888; Andor, UK). The z-focus was controlled by a commercial perfect focus system (Nikon, Germany). Acquisitions were controlled by μManager (Edelstein et al., 2014).

### Total internal reflection fluorescence (TIRF) microscopy setup

The TIRF microscope used for fluorescent protein evaluation was a commercial setup, VisiScope (Visitron Systems, Germany), consisting of a Nikon Ti Eclipse microscope body equipped with an CFI Apo TIRF 100x oil objective (NA 1.49, Nikon). Laser illumination coupling (405 nm and 561 nm lasers in the Visitron TIRF module) was achieved via optical fibers. Excitation and emission were separated and filtered by the ET - 405/488/561/640nm Laser Quad Band Set (Chroma, USA). Fluorescence was detected by an emCCD (iXON Ultra 888; Andor, UK). The z-focus was controlled by a commercial perfect focus system (Nikon, Germany). The setup and data acquisition were controlled with the Visiview 3.0.1 software (Visitron Systems, Germany).

### Evaluation of fluorescent protein performance

The measurements were performed on living WR806 cells immobilized on agarose pads, which expressed cytosolic fluorescent proteins from pTA231 plasmids. First, cells were exposed to 561 nm light, to bleach all possible preactivated/preconverted fluorescent protein signal. After taking snapshots (561 nm laser, 200 W/cm^2^, 20 frames with 60 ms exposure), which were used to measure the average background signal in cells, regions of interest were exposed to 405 nm light (4.5 J/cm^2^) to photoconvert/photoactivate fluorescent proteins and another series of snapshots with the 561 nm laser was taken to measure the average fluorescence intensity. The ratio of the average signal per cell before and after photoactivation/-conversion is plotted in **Figure 1c**.

### Single-molecule imaging of FtsZ1 and RNA Polymerase

All single-molecule measurements were performed in a customized heating chamber set to 42°C. Strains were imaged using the following laser intensities, unless state otherwise: 405 nm 0.5- 15 W/cm^2^; 488 nm 1-10 W/cm^2^; 561 nm 800 W/cm^2^; 730 nm 450 W/cm^2^. 405 and 488 nm lasers were pulsed every 10^th^ frame and their intensities were gradually increased during imaging to keep the density of photoactivated/-converted signal at a constant level. For each field of view (FOV), 10.000 frames were collected at 16.67 and 33.3 Hz for imaging the FtsZ1 and RNA Polymerase proteins, respectively.

### Dual-color imaging of cells expressing FtsZ1-Dendra2Hfx and FtsZ1-PAmCherry1Hfx

Cultures with strains expressing fluorescently labeled FtsZ1 at the same OD level of 0.2 were mixed together and loaded on an agarose pad. For comparison of the Dendra2Hfx photoconversion and PAmCherry1Hfx photoactivation efficiency using 405 nm light, FOVs containing both strains were imaged (16.67 Hz frame rate, 10000 frames) using either 0.5 W/cm^2^ or 1.5 W/cm^2^ 405 nm laser intensity, pulsed every 10^th^ frame. For the sequential dual-color readout, imaging protocols adapted from (Virant et al., 2017) were used. For FOVs containing both strains, first bright light images as well as images of green pre-converted Dendra2Hfx fluorescence were taken. Dendra2Hfx was read-out using primed photoconversion (488nm 1-10 W/cm^2^ and 730 nm 450 W/cm^2^). After the full readout of Dendra2Hfx, PAmCherry1Hfx was read-out by UV-photoactivation (405 nm 0.5-15 W/cm^2^).

### Data analysis

All cells were segmented based on their bright light images using Fiji (Schindelin et al., 2012). Segmented areas were used for the analysis of FP performance as well as for molecular counting and determination of cellular areas. The average cell fluorescence intensity quantification before and after photoactivation/- conversion was done in Fiji. Localizations of single fluorescent spots were obtained by rapidSTORM (Wolter et al., 2012) and tracked and analyzed by *swift* (Endesfelder et al.). Achieved experimental single-molecule localization precisions were measured by the NeNA approach (Endesfelder et al., 2014).

Molecular counting of FtsZ1 and RNA polymerase molecules per cell and cell area analysis was done with a custom script written in Python 3.7 with NumPy, Pandas and Tifffile libraries. FtsZ1 ring thickness analysis was performed with a self-written script in Fiji. All box charts and histograms were created using Origin (OriginLab) and figures were assembled using Adobe Illustrator (Adobe Inc.). In the histograms of **Figure 4**, weighted mean jump distances are plotted. This way, each single-molecule trajectory contributes to the histogram proportionally to its number of steps.

## Results

### Establishing SMLM imaging in *Haloferax volcanii*

*Haloferax volcanii* poses two major technical challenges for single-molecule fluorescence microscopy imaging. First, they naturally synthesize a high level of carotenoid pigments such as bacterioruberin, which cause some autofluorescence background signal. This heavily affects SMLM imaging, as SMLM is based on the detection of individual fluorophores (**Supplementary Text**). Secondly, as obligate halophilic extremophiles, they require growth under high salt concentration and grow best at elevated temperature (42-45°C). To establish SMLM imaging for *H. volcanii* as shown in **Figure 1a** for the cell division protein FtsZ1, we have addressed both of these issues.

From previous reports it is known that the deletion of the gene HVO_2528, which encodes for the *crtl* phytoene dehydrogenase, interrupts carotenoid synthesis and results in colorless *H. volcanii* cells without showing additional effects on phenotype or growth (Stachler et al., 2017;Maurer et al., 2018). We created the stable deletion-mutant strain WR806 and compared it with the red-colored *H. volcanii* laboratory strain H119 which synthesizes carotenoids (**Figure 1b**). Cultured in colorless Hv-Cab medium (Duggin et al., 2015;de Silva et al., 2020), WR806 appeared white on agarose plates and hardly distinguishable from background noise under SMLM imaging conditions, while H119 colonies were red-colored on plates and the cells exhibited an autofluorescence signal. This and the fact that the deletion did not cause any changes in cell morphology (**Supplementary Figure 1**) supports the choice of WR806 as host strain for SMLM imaging.

Next, we assessed which SMLM-compatible fluorescent proteins (FPs) could be potentially used as genetic fluorescent labels in *H. volcanii*. We selected three well-performing FPs derived from natural FPs of different biological origin: photoactivatable mCherry1 (PAmCherry1), derived from sea anemone Discosoma sp. (Subach et al., 2009), and the two photoconvertible proteins Dendra2 from octocoral Dendronephthya sp. (Gurskaya et al., 2006), and mMaple3 from Clavularia sp. (Wang et al., 2014). We expressed each FP from backbone pTA231 to the cytosol of WR806 host cells and evaluated their expression and functionality by measuring the cellular signal before and after photoactivation/-conversion of the FPs by UV light (**Figure 1c**). While cells expressing mMaple3 showed no substantial signal and the cells were barely distinguishable from WR806 control cells (not expressing any fluorophore), a signal increase was observed for cells expressing PAmCherry1 (about 5-fold) and for Dendra2 (about 2-fold). Additionally, we did not observe any change in cellular morphology when compared to H119 and WR806 control strains (**Supplementary Figure 1**). In contrast, cells expressing mMaple3 were elongated and rod-shaped. Consequently, we codon-optimized PAmCherry1 and Dendra2 sequences but did not further investigate the reasons for absent fluorescence signal and altered morphology of mMaple3 expressing cells.

We then evaluated codon-optimized versions of PAmCherry1 and Dendra2 for expression in *H. volcanii*. For both, PAmCherry1Hfx and Dendra2Hfx, the fluorescence signal in cells increased significantly (approx. 20-fold) (**Figure 1c**) which confirms efficient expression. Also, cell morphology remained unchanged (**Supplementary Figure 1**). Both, Dendra2 and Dendra2Hfx showed a more consistent read-out within experiments (smaller spread of individual cellular signals) as well as for different independent replicates of experiments (color-coded in **Figure 1c**, the mean of each individual experiment is marked by a diamond symbol), when compared to PAmCherry1 and PAmCherry1Hfx. We did not evaluate this finding further, but it suggests that Dendra2 might be a more suitable choice when choosing a genetic tag for quantitative SMLM studies (**Figure 1c**).

### Benchmarking single- and dual-color SMLM imaging in *Haloferax volcanii*

Using WR806 as host and Dendra2Hfx and PAmCherry1Hfx as suitable fluorescent labels, we benchmarked our SMLM imaging strategy on the division ring protein FtsZ1, which has been studied using immunofluorescence and GFP-fusions in *H. volcanii* (Poplawski et al., 2000;Duggin et al., 2015;Walsh et al., 2019). FtsZ1 was C-terminally tagged with either PAmCherry1Hfx or Dendra2Hfx and expressed in WR806 cells from a pTA962 backbone, which was previously used for FtsZ1-GFP studies (Duggin et al., 2015). FtsZ1 ring structures were clearly visible for all constructs as shown in **Figure 2a**. For both, PamCherry1Hfx and Dendra2Hfx, FtsZ1 structures were present at mid cell and could be resolved at a spatial resolution of about 24 nm. This revealed structural details such as branching rings and apparent double ring or helical arrangements.

We then compared the SMLM image quality of FtsZ1-PAmCherry1Hfx and FtsZ1-Dendra2Hfx structures. Cultures of WR806 cells expressing FtsZ1-PAmCherry1Hfx and FtsZ1-Dendra2Hfx were mixed together and imaged under several different UV nm light intensities (**Supplementary Figure 2**). The strains can be easily differentiated in the green channel as pre-activated PAmCherry1Hfx remains dark while Dendra2Hfx displays green, pre-converted GFP-like fluorescence. Whereas an UV light intensity of 0.5 W/cm^2^ was sufficient to efficiently record SMLM images of FtsZ1-Dendra2Hfx, FtsZ1-PAmCherry1Hfx constructs remained almost completely dark (**Supplementary Figure 2a**, upper panel and **Supplementary Figure 2b**). Increasing the UV intensity led to some photoactivation of FtsZ1-PAmCherry1Hfx (**Supplementary Figure 2a**, lower panel and **Supplementary Figure 2b**), but similar performance as for FtsZ1-Dendra2Hfx was only achieved at an about 20-fold increased intensity of 10 W/cm^2^ (**Supplementary Figure 2b**). We also compared the brightness of individual fluorescent spots. Dendra2Hfx signal was slightly brighter than PAmCherry1Hfx (251 versus 217 photons per fluorescent signal, median values, **Supplementary Figure 2c**). These results suggest that Dendra2Hfx is superior to PAmCherry1Hfx and can, due a 20-fold lower UV-photoconversion threshold intensity, minimize phototoxicity effects in SMLM studies.

In addition to UV-photoconversion, Dendra2-like proteins possess the ability to be photoconverted by an alternative photoconversion pathway, called primed photoconversion (PC), mediated by visible light (Dempsey et al., 2015;Turkowyd et al., 2017). Previously, we reported that PC works efficiently in a pH range of 7-10 and can be used for live-cell SMLM studies with reduced phototoxicity (Turkowyd et al., 2017). Testing PC on *H. volcanii* cells, we find that PC works efficiently and resulted in high quality SMLM images (**Figure 2a, right panel**). Thus, by choosing Dendra2Hfx, UV-photoconversion can successfully be replaced by PC, reducing the potential phototoxicity even further. Even though the pH of the cytosol of *H. volcanii* cells remains unknown, we can estimate from our efficient PC rate that it is based around pH 7-10, the optimal range for PC.

We previously also demonstrated that PC can be combined with UV-photoactivation into a quantitative dual-color SMLM imaging mode (Virant et al., 2017). To test this strategy for *H. volcanii*, we again mixed WR806 cells expressing either FtsZ1-PAmCherry1Hfx or FtsZ1-Dendra2Hfx (**Figure 2b**). We then used PC to image FtsZ1-Dendra2Hfx and observed that FtsZ1-PAmCherry1Hfx cells remained dark, keeping the FP in its pre-activated form (**Figure 2b I**.**a and II**.**a**). After the full read-out of Dendra2Hfx, we imaged FtsZ1-PAmCherry1Hfx using UV-photoactivation. We then detected only FtsZ1-PAmCherry1Hfx signal but no residual FtsZ1-Dendra2Hfx (**Figure 2b I**.**b** and **II**.**b**). The combination of PC- and UV-SMLM imaging therefore provides an elegant method for dual-color SMLM imaging in *H. volcanii* as it combines the high labeling specificity and efficiency of two genetic labels (and as such circumvents typical staining problems when using extrinsic SMLM dyes) with the aberration-free and quantitative read-out in one SMLM imaging channel. Future biological studies on two different proteins of interest within the same cell will be reliably recorded in high SMLM quality when using this strategy.

Finally, we tested whether is it possible to co-stain chromosomal DNA and evaluated DNA stains that were successfully used in live *H. volcanii* (Samson et al., 2008;Delmas et al., 2013). Acridine orange emits fluorescence close to most SMLM-compatible FPs, which limits its use as a co-stain to complicated protocols where the dye is only added after SMLM imaging. DAPI, Hoechst 33258 and Hoechst 33342 are excited using UV-light, which photoactivates/-converts the FP labels. However, as Dendra2 can be converted using PC, it can be imaged without photobleaching UV-excitable co-stains, and the chromosomal DNA can be visualized afterwards. **Supplementary Figure 3** shows exemplary cells where FtsZ1-Dendra2Hfx was co-imaged with DNA using Hoechst 33342.

### Imaging subcellular organization and dynamics of proteins in living *H. volcanii*

After having SMLM imaging established for *H. volcanii*, we aimed at imaging exemplary fusion proteins in greater detail. We chose two examples, the in Figure 1 and 2 already showcased division protein FtsZ1, and the highly dynamic RNA polymerase, responsible for transcription. For both, we selected Dendra2Hfx as label. To demonstrate how SMLM imaging can help studying archaeal cell biology, we decided for the practical example of imaging the cells under different growth conditions and cultivated our strains either using the more limiting Hv-Cab medium or the rich Hv-YPC medium. Motivated by bacterial SMLM studies in *Escherichia coli*, where the spatial organization of RNA polymerases was shown to be highly influenced by growth conditions (Endesfelder et al., 2013), we expected that differences in cellular metabolism should influence the spatiotemporal organization of both proteins.

### Resolving the FtsZ1-Dendra2Hfx ring structure in *H. volcanii*

FtsZ1 data confirmed that the protein indeed forms diverse architectures and suggested that growth conditions have a substantial impact on the FtsZ1 organization (**Figure 3a**) as well as cell morphology (Duggin et al., 2015;de Silva et al., 2020). Cells growing in Hv-Cab medium (**Figure 3a, left**) have irregular or round shapes and most cells possess single rings at mid cell. In contrast, cells cultivated in YPC medium (**Figure 3a, right**) are more often rod-shaped (Delmas et al., 2013;Duggin et al., 2015;de Silva et al., 2020), their FtsZ1 structures appeared more diverse and FtsZ1 rings are slightly thicker.

Next, we quantified these observations (**Figure 3b** and **3c**). Despite different morphologies in both conditions, cells comprised similar cellular areas (**Figure 3b**). The number of labeled FtsZ1 molecules per cell (including those not involved in the ring formation but that are diffusive in the cytosol) in cells growing in YPC is similar to those in Hv-Cab with median values of 213 and 330 copies, respectively (**Figure 3b**). Importantly, as FtsZ1-Dendra2Hfx was expressed transiently from a plasmid this does not represent the full number of FtsZ1 molecules in the cell. Here, we do not know the number of native, unlabeled FtsZ1. A study that aims to quantify FtsZ1 copy numbers would need a stable cell line expressing FtsZ1-Dendra2 from its native locus.

The ring thickness as measured for pre-converted green Dendra2Hfx as well as for photoconverted Dendra2Hfx in the SMLM image differed significantly due to the considerably improved resolution of SMLM data. In epifluorescence, the ring structures are convoluted due to the diffraction-limited signal and appear to be hundreds of nanometers thick. The data suggests only a slight difference in ring thicknesses between both media. For SMLM data, their difference becomes apparent: For cells grown in YPC, rings are about 2.5 times thicker with mean values of 179 nm and 70 nm for YPC and Hv-Cab, respectively (**Figure 3b**). Again, we cannot access the native, unmarked FtsZ1, but argue that this does not affect the above results. We assume that FtsZ1 and FtsZ1-DendraHfx are completely mixed in the structures and are not located in separate subareas.

Finally, we classified the apparent FtsZ1 structures in five groups: full, branching, double, helical and incomplete rings (**Figure 3c**). While our diffraction-limited data with pre-converted Dendra2Hfx reproduced well the findings of previous studies with mainly single FtsZ1 rings (Duggin et al., 2015;Walsh et al., 2019), we observed in the SMLM images of the same cells structures deviating from a single ring at a much higher rate. Especially for growth in YPC, the FtsZ1 architectures are highly diverse: only 30.8% of the cells have a full ring and another 24.4% show incomplete rings. The other cells showed more complex structures with 22.6% with branching rings, 11.3% with helices and 10.9% with apparent double rings. These first SMLM results thus provide an ideal basis for more detailed future studies to explore the molecular mechanisms of FtsZ1 and FtsZ2 ring formation in cell division (Liao et al., 2020).

### Single-molecule tracking of RNA polymerase in *H. volcanii*

Compared to FtsZ1, RNA polymerase is expected to be a more dynamic molecule, diffusing through the cytosol and scanning the DNA for promotor sequences to initialize transcription. Evaluating possible labeling strategies, we chose to label the RNA polymerase subunit RpoD. We performed experiments similar to our FtsZ1 experiments albeit at a faster imaging speed at a frame rate of 33 Hz to account for the faster dynamics of the protein. As for FtsZ1, we expressed RpoD-Dendra2Hfx transiently from a plasmid with expression of unlabeled RpoD from its native locus still present. In this imaging mode, we could follow the dynamics of individual RNA polymerase molecules in cells grown in either Hv-Cab or YPC medium, and took a DNA snapshot using Hoechst 33342 to map the spatial co-organization of RNA polymerases and chromosomes (**Figure 4a**). We tracked the fluorescent signals of each molecule and reconstructed cellular diffusion maps. We then inferred the motion type(s) for each molecule and categorized them into immobile and diffusive fractions, which can be attributed to actively transcribing (immobilized on DNA) and diffusing DNA-scanning RNA polymerases (**Figure 4a**, red and violet trajectories, respectively), as shown before in bacterial studies for *Escherichia coli* (Bakshi et al., 2013;Stracy et al., 2015). **Figure 4b** shows exemplary cells for both growth conditions where the classified trajectories are overlaid onto the DNA image. We observed a substantial difference in cell sizes between the different media, which we, to that extent, had not observed before for the strains expressing FtsZ1-Dendra2Hfx from the same backbone pTA962 under the same growth conditions. When expressing RNA-polymerase from pTA962, cells growing in rich YPC medium have almost 2-fold larger area in comparison to cells growing in Hv-Cab (**Figure 4b**). Furthermore, cellular morphology remained round for RNA polymerase expressing cells.

For some cells, we observed only few trajectories in their mid-cell area, and especially immobile ones often located on cellular peripheries, yet not strictly in proximity to the cell membrane (red trajectories, **Figure 4b**). These findings are similar to observations made for *E. coli* cells (Stracy et al., 2015). We then quantified the percentages of transcribing and diffusive labeled RNA polymerases per cell (**Figure 4b**). Histograms presenting the distributions of weighted mean jump distances (for details see Methods) show only minor differences and 18%, respective 14% of single-molecule dynamics are assigned to immobile RNA polymerases for Hv-Cab and YPC media. This result holds true when counting labeled RNA polymerase molecules: we observed a median of 15 immobile labeled RNA polymerase molecules per cell in Hv-Cab (which is equivalent to 19% of in total 81 labeled RNA polymerase copies per cell) and 17 labeled immobile RNA polymerase molecules in YPC (which is equivalent to 15% of the total of 111 labeled RNA polymerase copies per cell) (**Figure 4b**). In summary, even though the overall number of labeled RNA polymerase copies is about 1.4 times higher in cells growing in full YPC media, the number of actively transcribing RNA polymerases remains relatively constant.

## Discussion

In this work, we have established protocols to study living *Haloferax volcanii* cells with quantitative SMLM methods. We created the carotenoid-free mutant strain WR806 to obtain autofluorescence-free *H. volcanii* cells and demonstrated that this strain is an excellent host for SMLM imaging. We further explored suitable SMLM-compatible fluorophores and demonstrated that Dendra2Hfx and PAmCherry1Hfx, codon-optimized FP versions for *H. volcanii*, work best. Dendra2Hfx performs better than PAmCherry1Hfx as it requires lower UV doses, is susceptible to PC and shows a higher brightness. Therefore, we recommend Dendra2Hfx as a first choice for SMLM studies when imaging a single target protein of interest. Nevertheless, PAmCherry1Hfx is also a suitable choice and can be used in addition to Dendra2Hfx in dual-color imaging by combining PC and UV-activation. Since Dendra2Hfx can be photoconverted without UV-light, a co-staining of the DNA by conventional UV-excitable DNA stains is also possible.

We then showcased the potential of SMLM imaging for investigating the cell biology of Haloarchaea using two biological test cases, the division protein FtsZ1 and the transcribing RNA polymerase, under different growth conditions. Our results on their variable copy number and spatiotemporal organization provide a strong incentive to engage in more detailed SMLM studies of the biology of both proteins and their interaction partners. This work therefore forms the basis for SMLM imaging as a well-established tool for investigating archaea and for exploring their unique biology in molecular detail in living cells in the near future. Building on the incentive of this work, we hope that SMLM strategies for other archaeal organisms, e.g. hyperthermophilic *Sulfolobus*, will eventually come within close reach.

## Conflict of Interest

The authors declare that the research was conducted in the absence of any commercial or financial relationships that could be construed as a potential conflict of interest.

## Author Contributions

UE and AM designed and supervised the study. ES and MM generated and validated the WR806 strain. JW and SS generated and validated all fluorescent constructs. JW, SS, BT, ID, UE and AM designed the individual experiments and JW, SS and BT performed and analyzed all experiments. BT and UE wrote the manuscript with the help of all authors.

## Funding

Work in the Endesfelder laboratory was supported by the Max Planck Society, the DFG priority program SPP 2141 (En1171/1-1) and by the Fonds der Chemischen Industrie. Work in the Marchfelder laboratory was funded by the DFG (Ma1538/25-1) in the frame of the DFG priority program SPP2141. Iain Duggin was funded by an Australian Research Council Future Fellowship (FT160100010).

## Acknowledgments

We thank Irma Merdian for expert technical assistance, Dina Grohmann for discussions and Uri Gophna for critical reading of the manuscript.

---

**Table 1.**
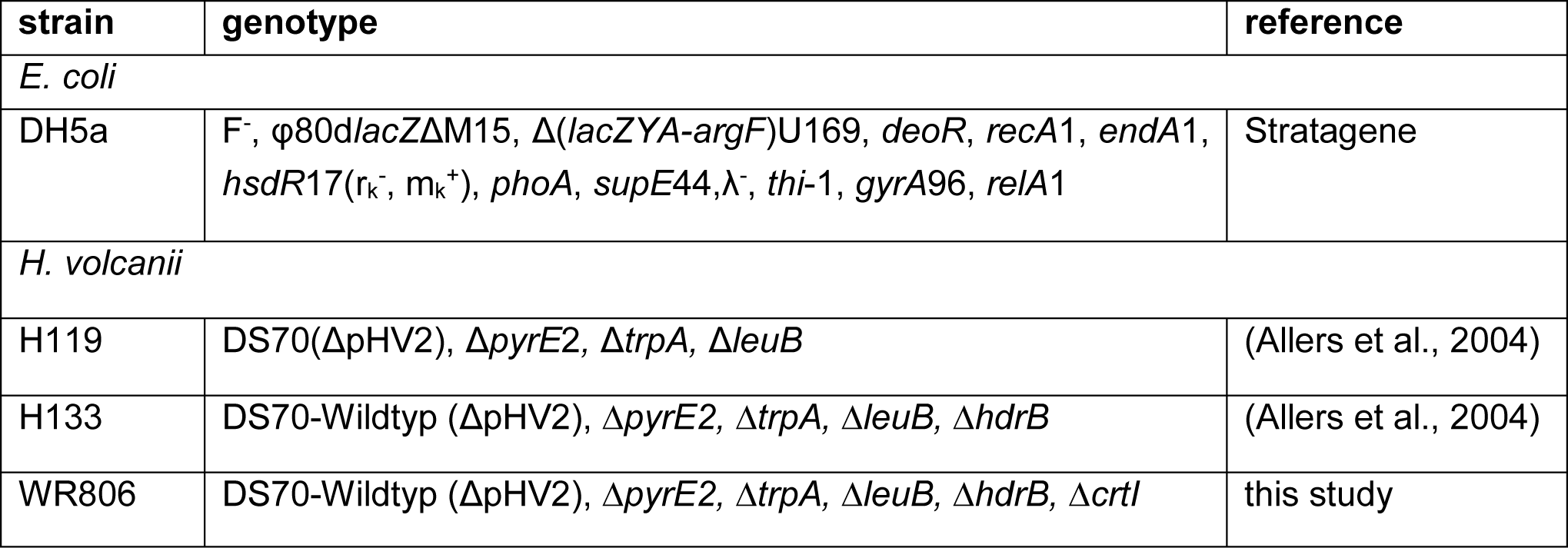
Strains used.

**Table 2.**
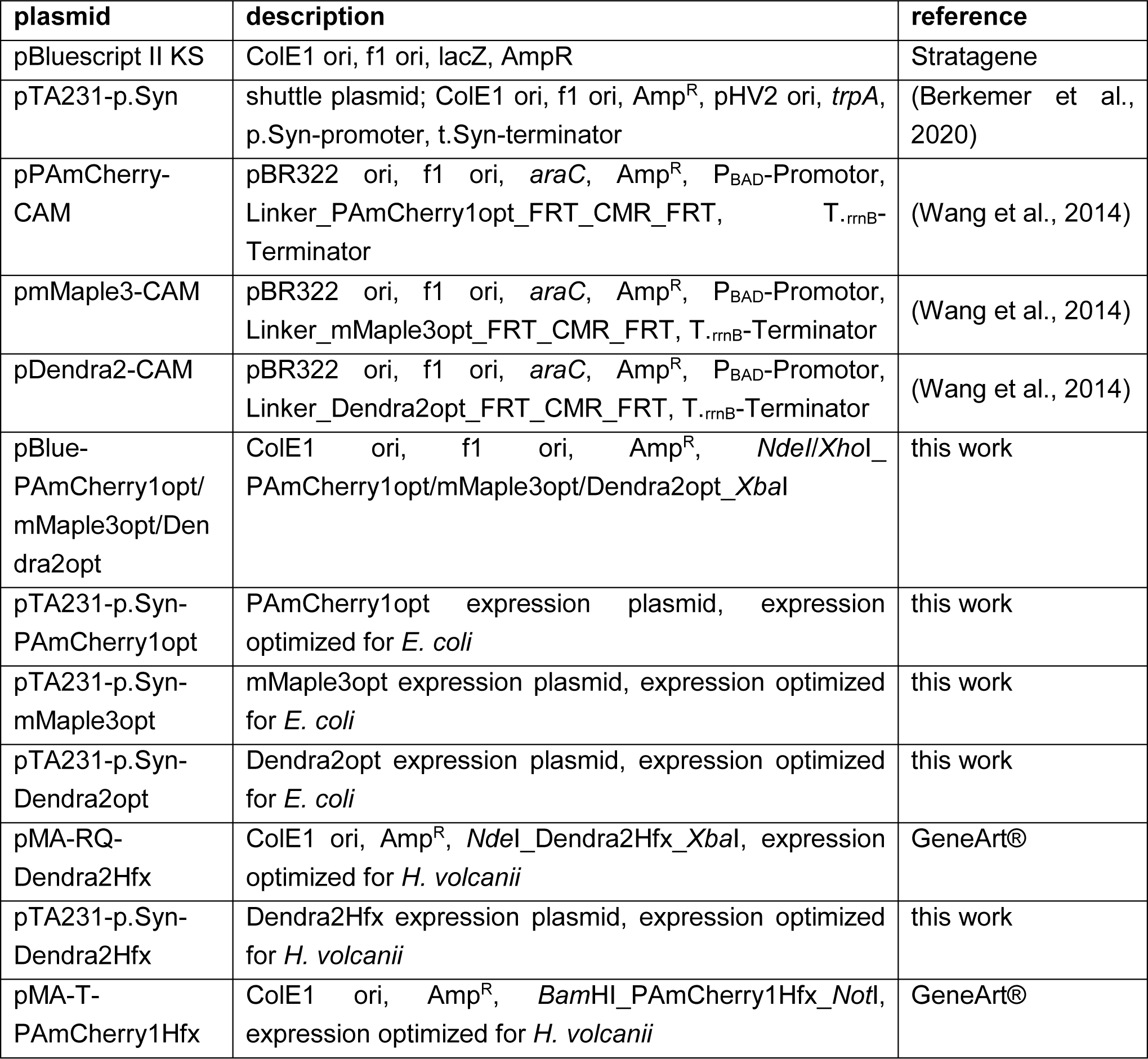
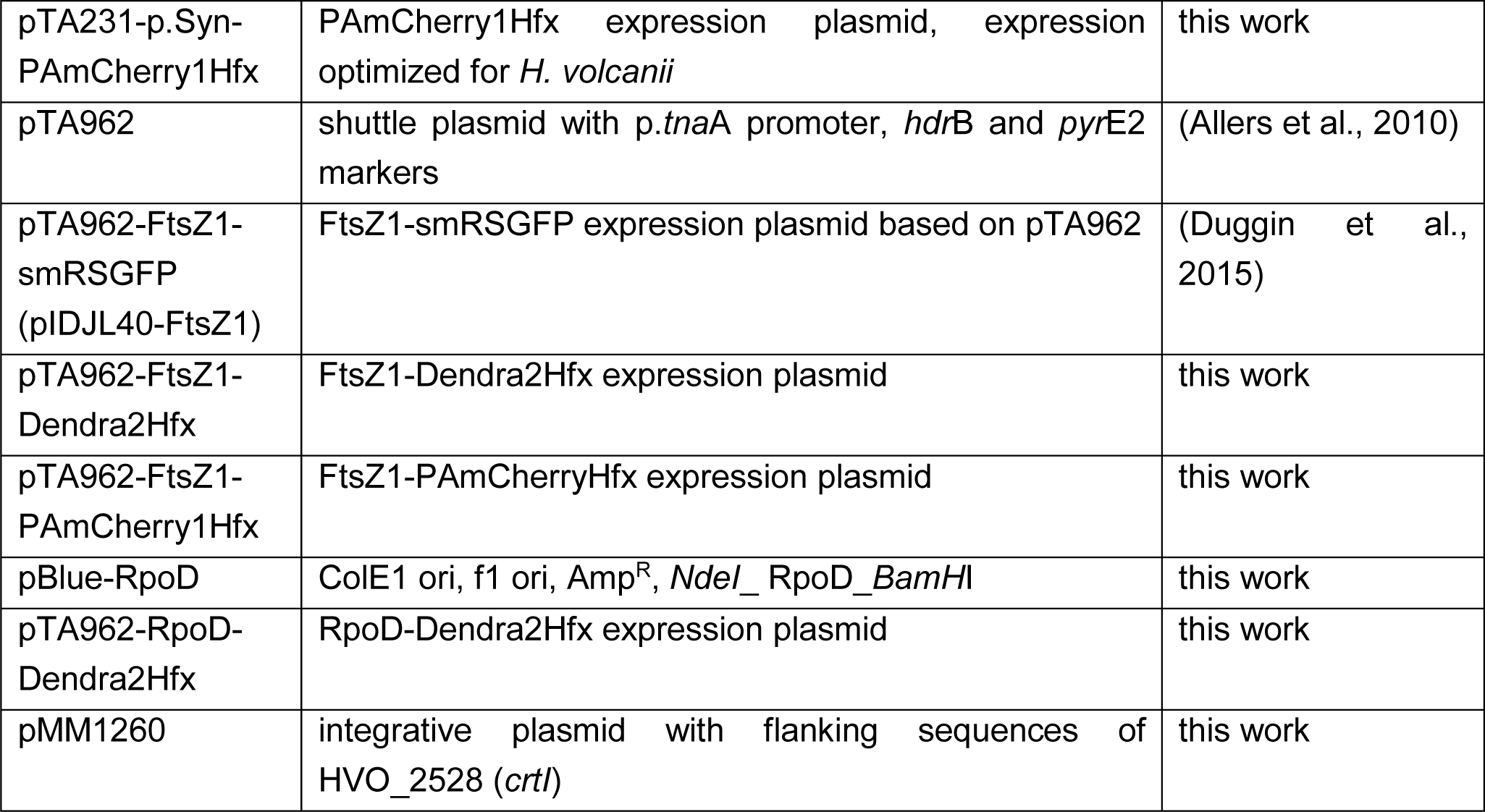
Plasmids used.

**Table 3.**
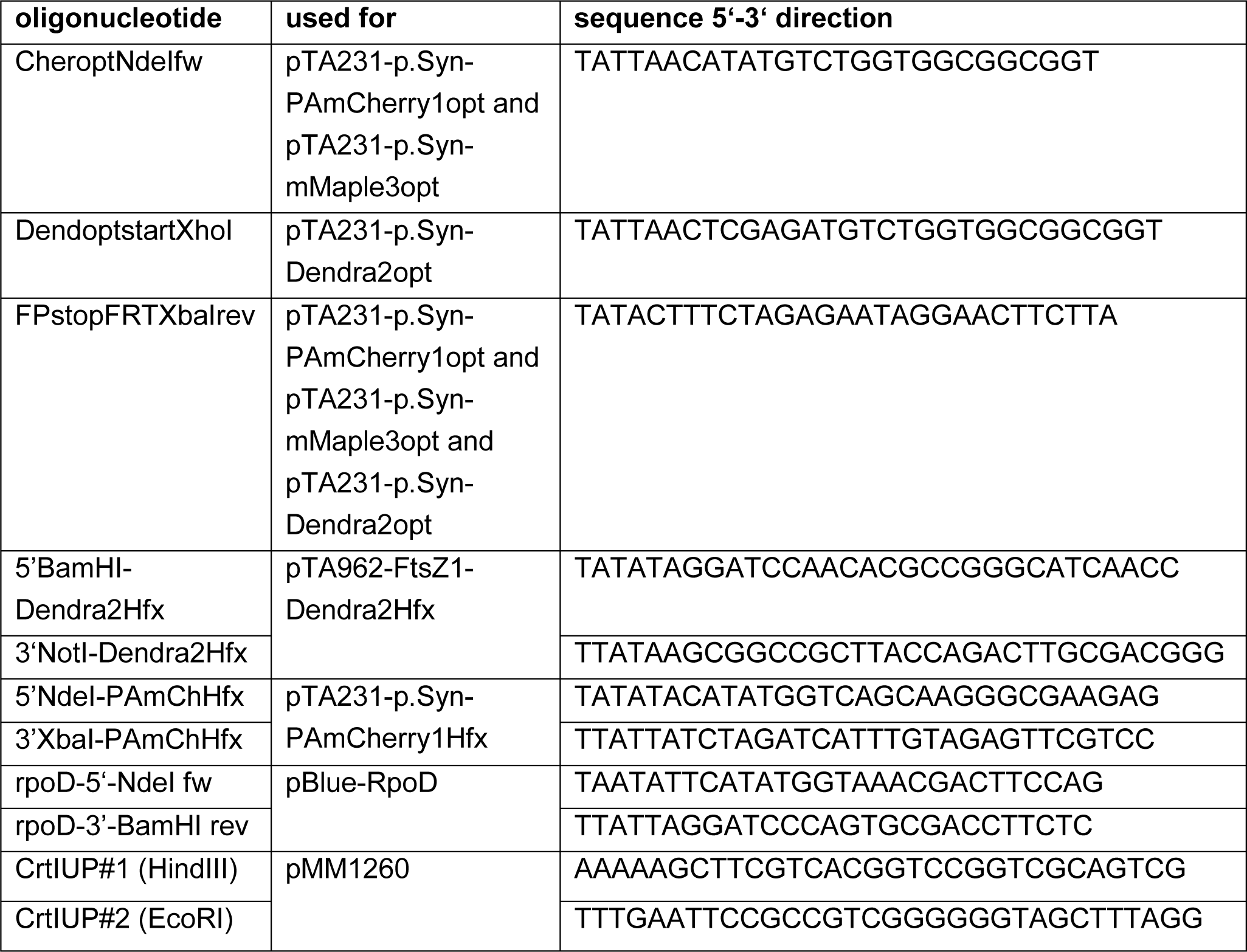
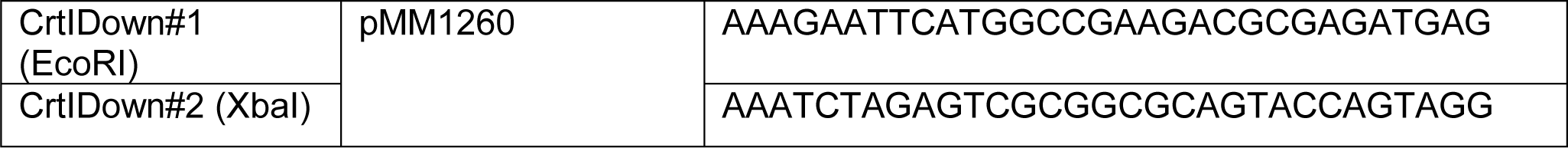
Oligonucleotides used.

## Notes

### Competing Interest Statement

The authors have declared no competing interest.

