## Supplementary Information for "Establishing live-cell single-molecule localization microscopy imaging and single-particle tracking in the archaeon *Haloferax volcanii*"

- **Supplementary Text**
- **Supplementary Figure 1**
- **Supplementary Figure 2**
- **Supplementary Figure 3**

### Supplementary Text

#### Background sensitivity of single-molecule imaging

Fluorescence microscopy studies can be compromised by autofluorescence background. While conventional fluorescence microscopy in wild-type red Haloferax cells is generally possible as already a few GFP together provide sufficient signal above background, single-molecule imaging is severely compromised, especially at high frame rates of dynamic imaging.

The background level is an important factor in the detection and localization of single molecules. If signals are not sufficiently brighter than the background, a large proportion of the weaker signals will remain undetected, and brighter signals that are still visible above the background are localized at lower precision, directly limiting the overall resolution of the super-resolved image.

In our measurements, WR806 cells significantly improve single-molecule imaging conditions for *Haloferax*: **Supplementary Figure 2** shows that our fluorescent proteins yield on average ~ 200-250 photons per image (corresponding to 21.000 to 27.000 AD counts), typical numbers for single-molecule imaging. The pixel size is optimized for the detection and localization of single fluorescent spots and individual spots are measured on an area of about 3x3 pixels. Thus, the average signal of a single fluorescent protein is not much stronger than the average background of 1.300 AD counts per pixel in H119 cells (**Figure 1b**), which results a 9-pixel background signal of 11.700 AD counts (translates into ~ 107 photons for our setup). In contrast, WR806 cells are indistinguishable from the background outside cells and single fluorescent protein signals can be detected above background.

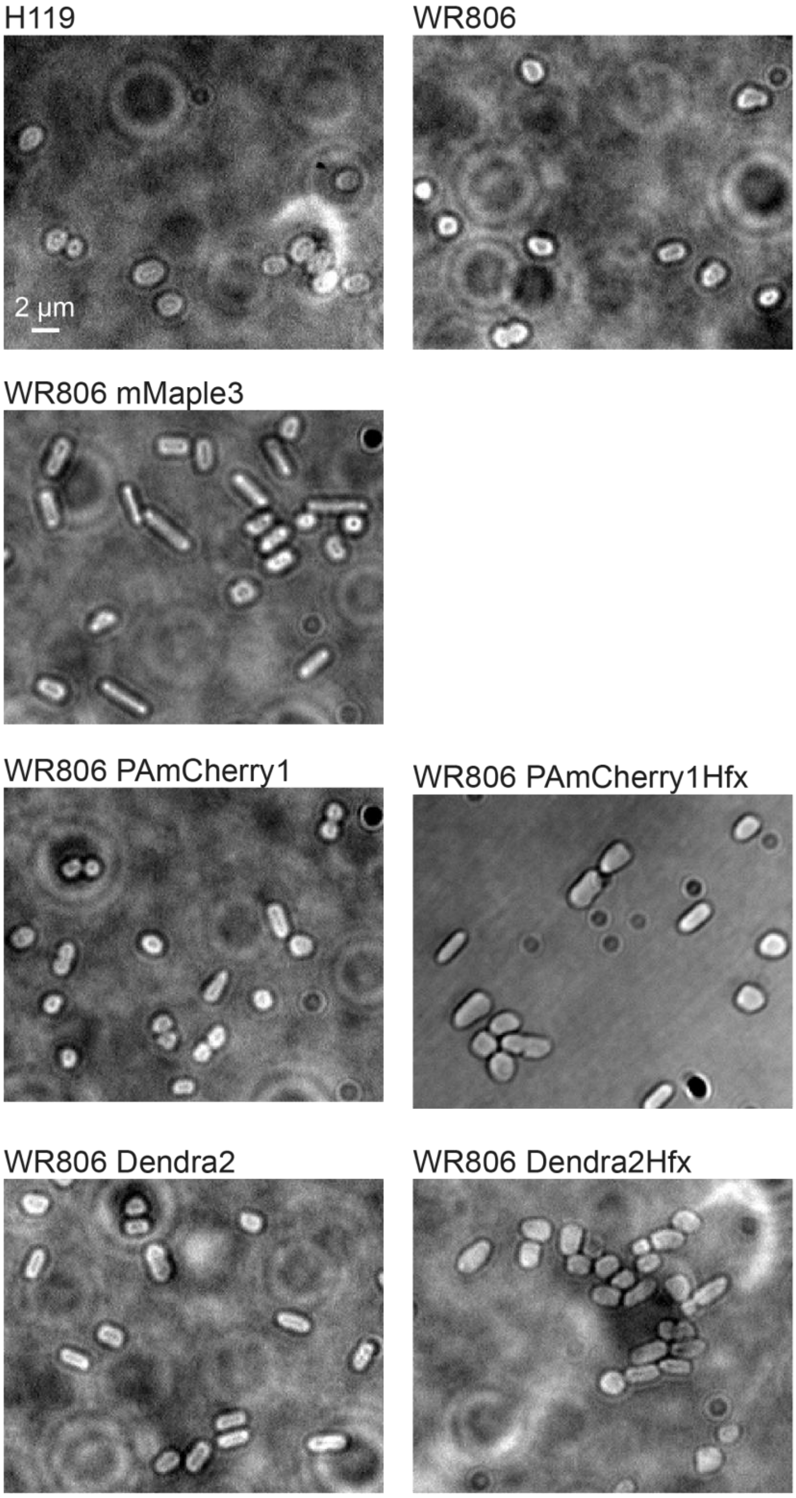


**Supplementary Figure 1. Cell phenotype varies under expression of different fluorescent proteins**

All strains expressing fluorescent proteins which showed efficient photoactivation/-conversion (**Figure 1c**) showed similar phenotypes which are comparable to the parental strains H119 and WR806. In contrast, the strain expressing mMaple3 shows strongly altered morphology. This and the fact that mMaple3 showed no fluorescence (**Figure 1c**) suggests improper protein folding that might cause increased cytotoxicity leading to a different growth phenotype.

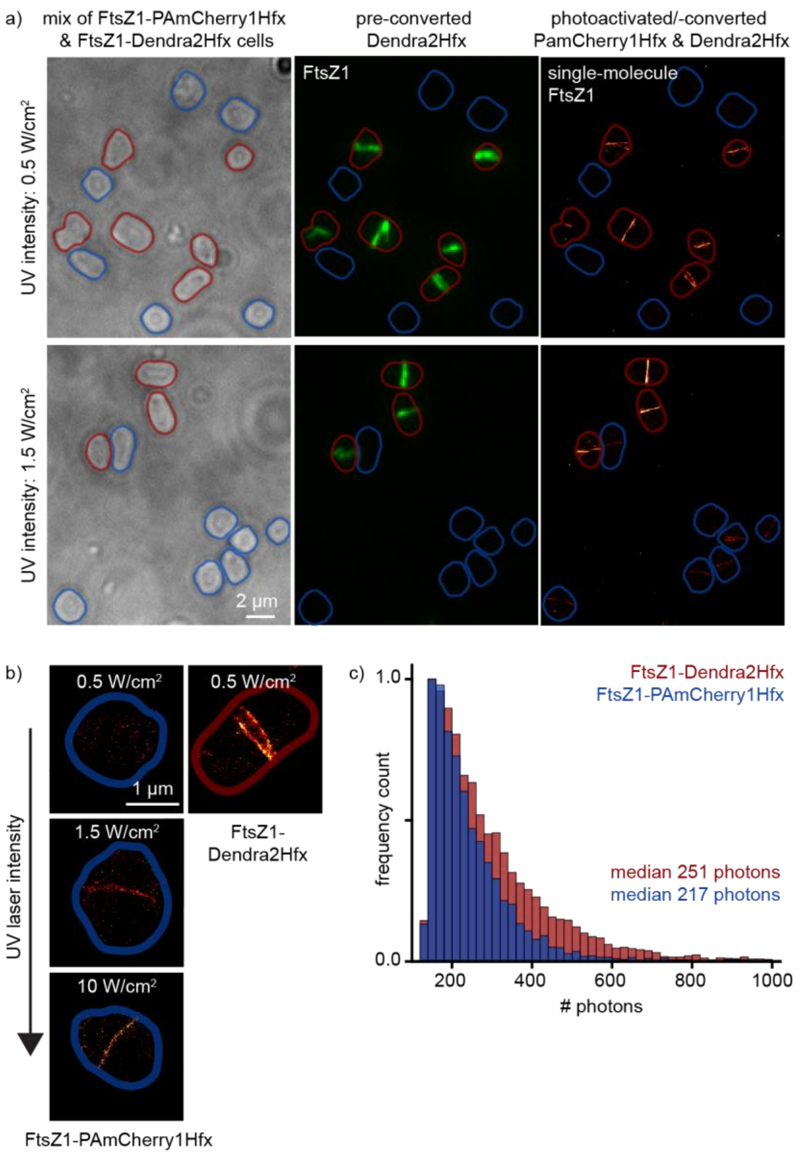


**Supplementary Figure 2. Dendra2Hfx performs superior to PAmCherry1Hfx.**
(a) WR806 cells expressing either FtsZ1-Dendra2Hfx or FtsZ1-PAmCherry1Hfx were mixed on the same agarose pad and imaged under different UV laser intensities. When 0.5 W/cm² of the UV light was applied, FtsZ1-Dendra2Hfx revealed good photoconversion which results in dense SMLM ring structures. In contrast, FtsZ1-PAmCherry1Hfx was not photoactivated. A read-out of FtsZ1-PAmCherry1Hfx became only possible when increasing the UV light intensity to 1.5 W/cm².
(b) To provide an image quality comparable to FtsZ1-Dendra2Hfx when imaging FtsZ1-PAmCherry1Hfx, the UV light intensity had to be significantly increased. 10 W/cm² was barely sufficient to achieve the same quality as for the FtsZ1-Dendra2Hfx structure when applying 0.5W/cm².
(c) Under imaging conditions as in (a), Dendra2Hfx emits on average more photons (median of 251 photons) than PAmCherry1Hfx (median of 217 photons) per fluorescent emission in a single imaging frame.

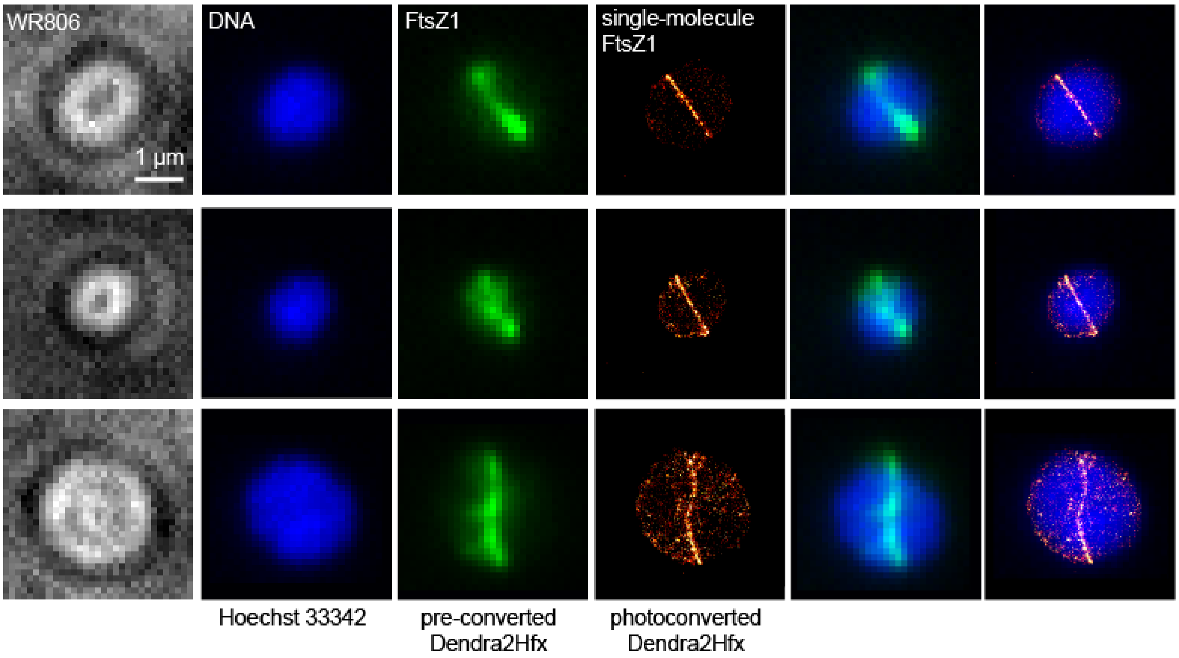


**Supplementary figure 3. Dual color imaging of FtsZ1-Dendra2Hfx and DNA stained with Hoechst 33342.** As FtsZ1-Dendra2Hfx can be photoconverted by primed photoconversion, which does not rely on UV light illumination, it is possible to use the DNA-intercalating dye Hoechst 33342 as a co-stain to obtain a DNA-reference image.
